# Usutu virus and West Nile virus use a transcellular route of neuroinvasion across an *in vitro* model of the human blood-brain barrier

**DOI:** 10.1101/2024.01.26.577380

**Authors:** Eleanor M. Marshall, Marion Koopmans, Barry Rockx

## Abstract

West Nile virus (WNV) leads to thousands of cases of severe neurological disease in humans each year. Usutu virus (USUV) is closely related to WNV, but rarely induces disease in humans. We hypothesised that USUV is less able to cross the blood-brain barrier, and is therefore less likely to infect the brain. Therefore, we developed an *in vitro* BBB model consisting of primary human brain microvascular endothelial cells (BMECs), pericytes and astrocytes. Both USUV and WNV invaded across the *in vitro* BBB via a transcellular mechanism in absence of barrier disruption. USUV replicated to lower titres than WNV but induced a comparable cytokine and chemokine response, with modulation of key factors associated with barrier function and immune-cell migration. In conclusion, USUV appears attenuated in its ability to replicate at this interface compared with WNV, but further work must to done to identify key determinants underlying the differing clinical presentations.

## INTRODUCTION

Many arthropod-borne viruses (arboviruses) can cause severe neurological disease in humans, but how these viruses bypass the numerous physical and immunological barriers in place to protect the central nervous system (CNS), is not well understood. West Nile virus (WNV) is a mosquito-borne flavivirus that causes thousands of cases of neuroinvasive disease in many regions of the world every year, including North America^1^, and increasingly in Europe^2^, with a continued emergence in new regions each transmission season. Usutu virus (USUV) is phylogenetically closely related to and co-circulates with WNV in Europe, but since the first case of human infection identified in 1981, there have only been around 100 documented cases of USUV induced disease and no reports of fatal infection^3^. WNV and USUV therefore appear to differ in their ability to cause severe disease in humans, which could stem from many factors, including the ability of the virus to gain access to the CNS via one or more routes of neuroinvasion. WNV is thought to use multiple routes of neuroinvasion, including both transcellular and paracellular invasion across the blood-brain barrier (BBB)^4^. The potential for use of these haematogenous routes of neuroinvasion by USUV is not well understood.

The BBB is a semipermeable selective border composed of tightly joined brain microvascular endothelial cells (BMECs), ensheathed by pericytes and astrocytes. This barrier is considered an important interface for invasion of viruses into the CNS from the blood. Such invasion can occur via a transcellular route in which the virus must infect or be transported across the cells of the barrier, or via a paracellular route in which virus is able to enter between the cells of a disrupted barrier, either as free virions or within infected immune cells in a so-called Trojan horse mechanism of invasion^4^.

Here, we aimed to determine whether USUV and WNV differ in their capacity to invade the CNS across the BBB. We employed a triple co-cultured *in vitro* transwell system, using primary human BMECs, astrocytes and pericytes, to recapitulate the human BBB and investigate the mechanism of viral invasion across this barrier. Identifying key differences and similarities between these two viruses will shed light on the essential virus or host factors that underlie viral neuroinvasion, and thereby aid in assessment of the future risk posed by USUV. We also provide a robust, biologically relevant platform that can be employed to evaluate the neuroinvasive capacity of emerging viruses and live-attenuated vaccine candidates, and to develop therapeutic strategies working to prevent viral neuroinvasion across the BBB.

## RESULTS

### USUV and WNV can infect and replicate within all three cell types of the human BBB

To identify the tropism of USUV and WNV for the different cell types of the human BBB we infected primary human BMECs, astrocytes and pericytes with USUV or WNV at an MOI of 1. Both USUV and WNV could infect and replicate in BMECs (**Fig. 1A**), with WNV peaking at 48hpi and showing a significantly higher titre over USUV at this time-point. WNV titres decreased by 72hpi, and did not show significant difference to USUV at this timepoint. Similar kinetics were observed in primary human astrocytes (**Fig. 1B**), with comparable titres for both viruses at 24hpi, leading to significantly higher WNV titres when compared with USUV at 48hpi. Both USUV and WNV showed decreasing in titres at 72hpi. In pericytes (**Fig. 1C**), USUV peaked at 24hpi, with a significantly higher titre than WNV at this timepoint. However, whilst USUV plateaued after 24hpi, WNV titres continued to increase, peaking at 48hpi.

**Figure 1.**
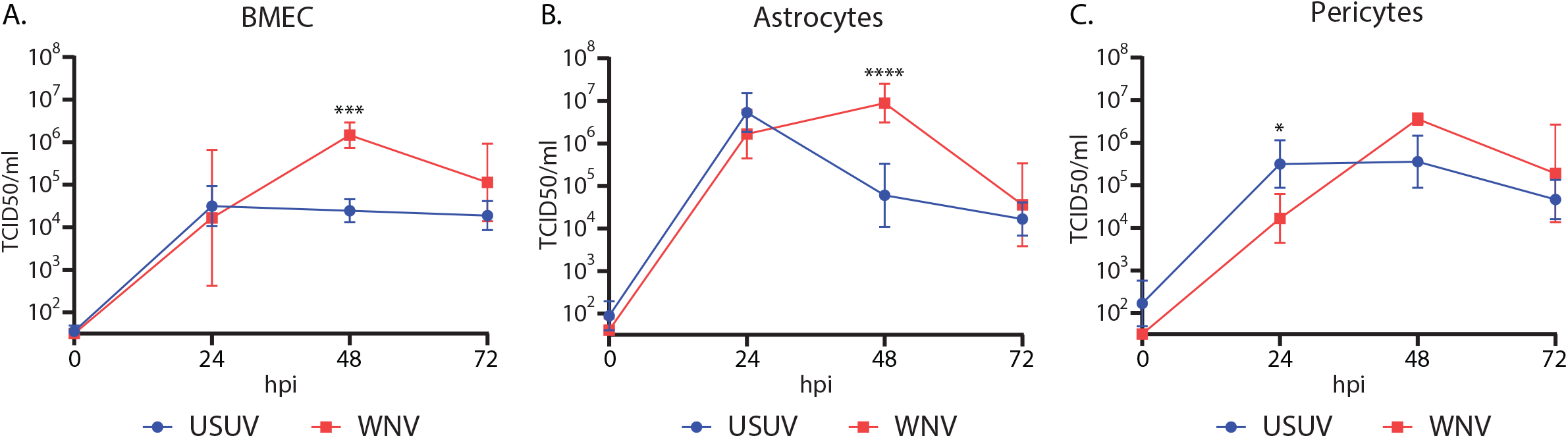
Usutu virus and West Nile virus can replicate in all cell types of the human blood-brain barrier. Replication kinetics of primary human A. brain microvascular endothelial cells (BMECs), B. astrocytes and C. pericytes infected with of Usutu virus (USUV) and West Nile virus (WNV) at a multiplicity of infection of 1. n=2. 3 replicates per condition, per experiment. * p =0.0158. *** p=0.0006. **** p<0.0001. Tukey’s multiple comparison.

We then carried out IF staining to confirm infection within the primary cell cultures, in combination with relevant cell markers. We identified staining for viral envelope of USUV and WNV in scattered foci throughout BMEC (**Fig. 2A**), astrocyte (**Fig. 2B**) and pericyte (**Fig. 2C**) cultures, and all three cultures stained for their expected markers. Overall, we found that all three cell types of the human BBB are susceptible to infection with USUV and WNV.

**Figure 2.**
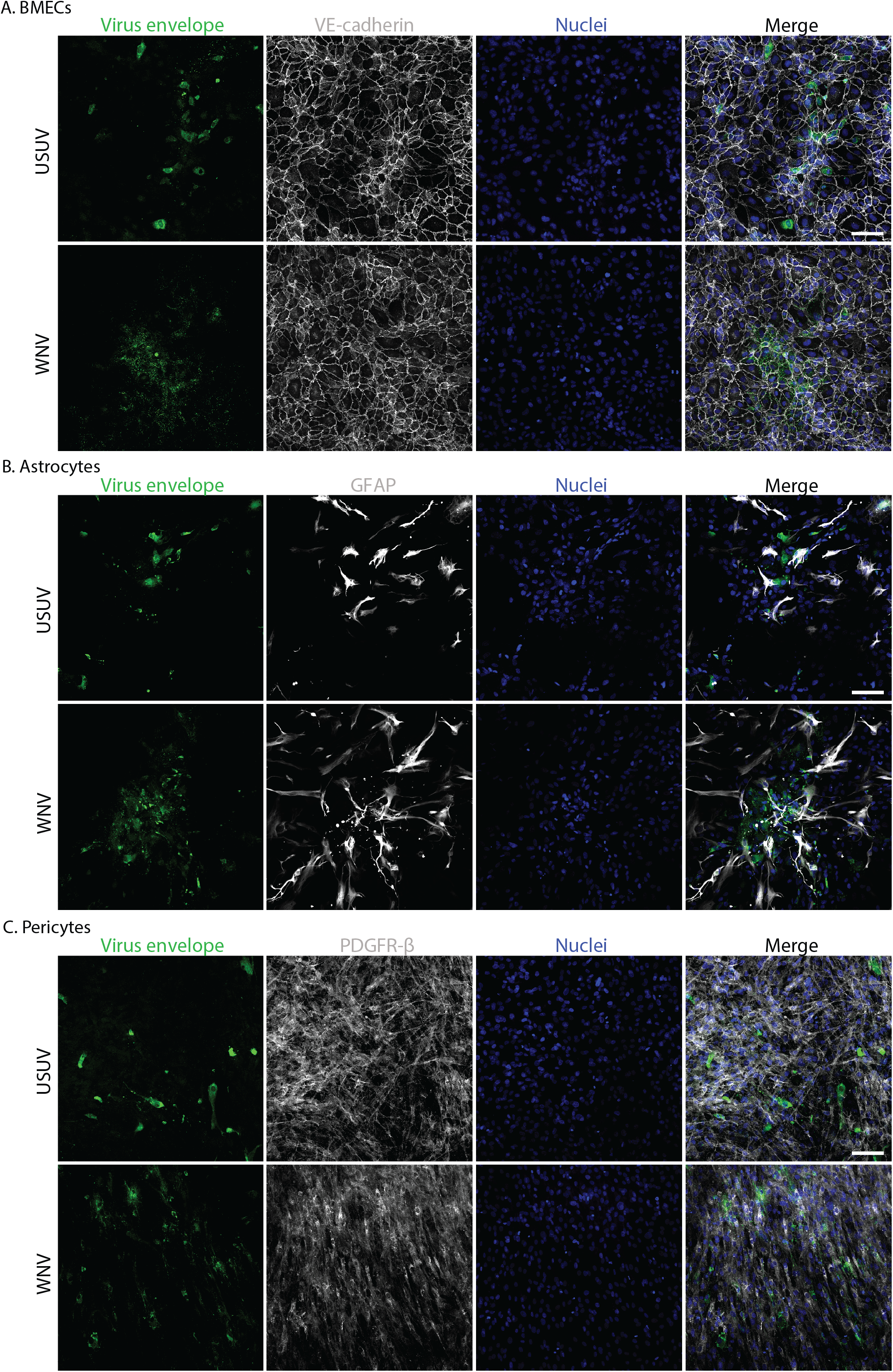
Usutu virus and West Nile virus infection can be visualised in all cell types of the human blood-brain barrier by immunofluorescent staining. **A.** Immunofluorescent (IF) staining of Usutu virus (USUV) and West Nile virus (WNV) infected brain microvascular endothelial cells (BMECs). VE-cadherin shown in white. **B.** IF staining of USUV and WNV infected astrocytes. GFAP shown in white. **C.** IF staining of USUV and WNV infected pericytes. PDGFR-β shown in white. Nuclei shown in blue. Virus envelope shown in green. Scale bars represent 100μm.

### The triple-cocultured *in vitro* blood-brain barrier exhibits the greatest barrier function

Next, to confirm data previously obtained in a similarly proximate co-culture model^5^, we investigated to what extent co-culturing the primary astrocytes and pericytes with BMECs contributed to barrier establishment. We seeded BMECs in the apical compartment alone, or in combination with either astrocytes, pericytes or both of these cell types seeded on the basolateral side of the transwell membrane (**Fig. 3A**). In absence of pericytes, astrocytes did not contribute to the barrier function, with the BMEC + astrocyte co-culture exhibiting low transendothelial electrical resistance (TEER) values similar to the monoculture of BMECs. The barrier of the BMEC monoculture peaked on day 7 post-culture establishment and the BMEC + astrocyte co-cultures peaked on day 6. The addition of pericytes did contribute to the barrier, with the BMEC + pericyte co-culture showing persistently higher TEER compared with the BMEC alone until day 8. The BMEC, astrocyte, pericyte triple co-culture showed the highest TEER, which rapidly increased between day 2 to 3 and stayed >2 fold higher than BMEC alone until day 6 post seeding when the triple-coculture dropped to TEER values similar to BMEC + pericytes (**Fig. 3B**). IF staining of the *in vitro* BBB confirmed the presence of all three cell types (**Fig. 3C**) with 3D rendering revealing the expected orientation in relation to the transwell membrane (**Fig. 3D, Vid. S1**). In addition to assessment of the barrier using TEER measurement, we also carried out IF staining of the BMECs in the apical compartment for the key tight-junction protein, ZO-1, and found that ZO-1 was expressed in our *in vitro* BBB system (**Fig. 3E**). These data suggest that co-culturing primary BMECs, astrocytes and pericytes has a synergistic effect on BBB barrier function.

**Figure 3.**
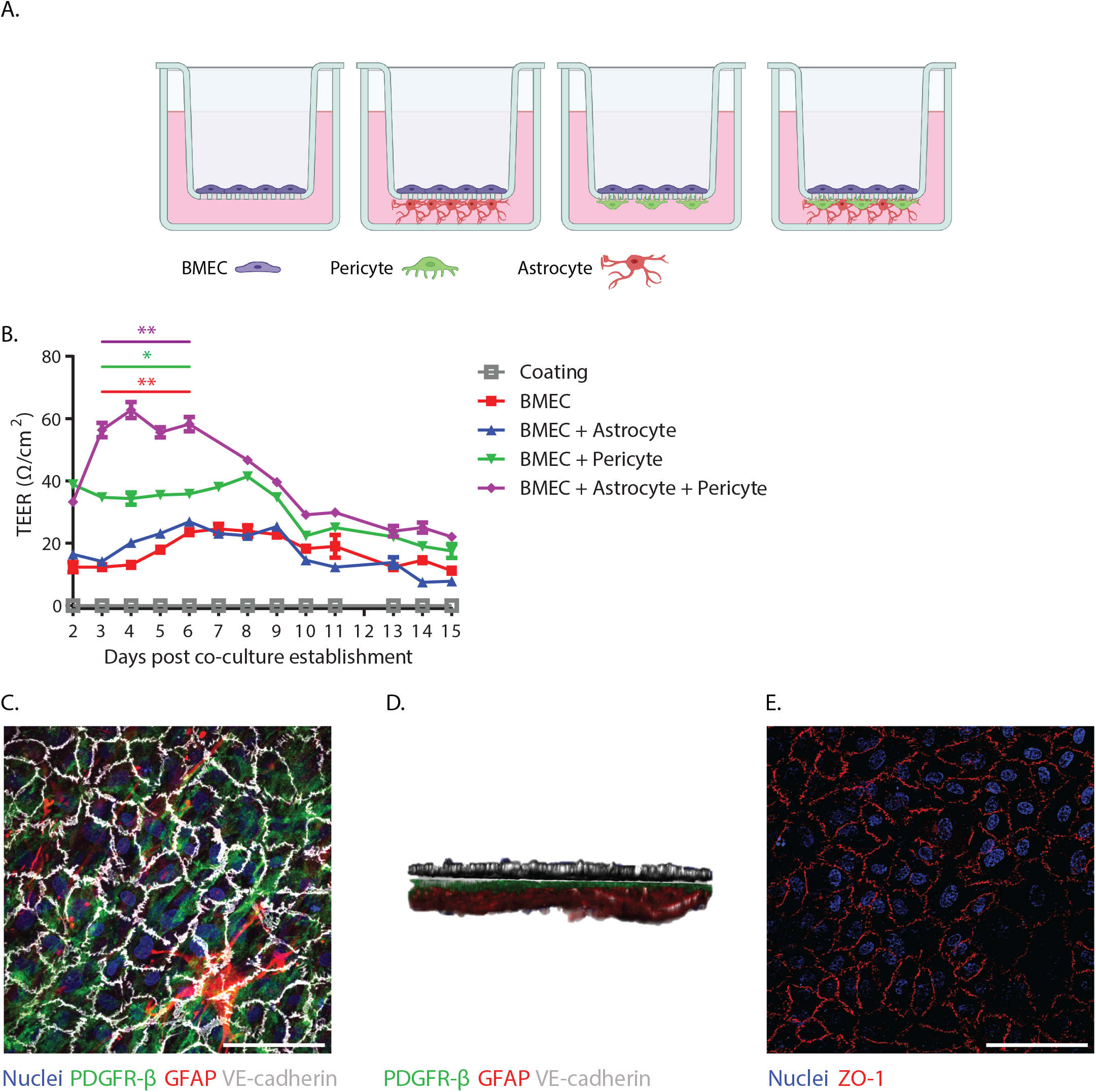
A triple-coculture of brain microvascular endothelial cells, astrocytes and pericytes provides the highest barrier function. **A.** Representation of the four different blood-brain barrier (BBB) culture set-ups, with brain microvascular endothelial cells (BMECs) in the apical compartment and astrocytes, pericytes or astrocytes and pericytes seeded on the basolateral membrane of the transwell. Made using biorender. **B.** Transendothelial resistance (TEER) of the BBB culture set-ups following culture establishment. 3 replicates per condition. Representative data from 3 independent experiments. Purple stars indicate BMEC monoculture vs BMEC + pericyte + astrocyte triple co-culture (** p<0.0051). Green stars indicate BMEC + pericyte double co-culture vs BMEC + pericyte + astrocyte triple co-culture (* p<0.0136). Red stars indicate BMEC monoculture vs BMEC + pericyte double co-culture (** p<0.0078). Tukey’s multiple comparison run on stated conditions for day 3 – day 6 post co-culture establishment. **C.** XY view of immunofluorescent (IF) stained triple-cocultured *in vitro* BBB viewed from the apical side. PDGFR-β shown in green. GFAP shown in red. VE-cadherin shown in white. **D.** XZ view of 3D rendered, IF stained triple-cocultured *in vitro* BBB. E. IF staining of ZO-1 in BMECs of apical compartment. Nuclei shown in blue. ZO-1 shown in red. Scale bars represent 100μm.

### USUV and WNV can invade across the *in vitro* BBB in absence of barrier disruption

To investigate early events of viral invasion, we initially performed a pilot with WNV to determine on what timescale initial invasion occurred. We found that viral genome could not be detected in the basolateral compartment until around 16hpi, assumedly after a first round of replication in BMECs (**Fig. S1A-D**). This lag was not caused by a blockade of virion diffusion by the coated transwell membrane (**Fig. S2**). Based upon this data, we chose both 16hpi and 24hpi timepoints to compare the kinetics of invasion for USUV and WNV, and also investigated whether USUV and WNV impact the barrier function on a longer timescale, following the initial invasion. As expected, WNV could not be detected in the basolateral compartment until the 16hpi time-point. USUV showed a similar pattern of invasion, increasing rapidly between 16hpi and 24hpi in the basolateral compartment. Titres in the apical compartment were comparable between USUV and WNV at 16 and 24hpi. USUV titres in the apical compartment peaked at 1.1×10^5^ TCID50/ml at 48hpi but WNV continued to increase to significantly higher titres compared with USUV at 48hpi and 72hpi, reaching titres of 2.9×10^7^ TCID50/ml. The basolateral compartments of each virus showed similar kinetics to the apical compartment, with WNV showing no significant difference between the compartments at any time-point. However, at 16hpi and 24hpi USUV did show a significant difference, with lower titres in the basolateral versus the apical compartment, but by 48hpi USUV titres were comparable in both compartments. (**Fig. 4A**). We confirmed infection of cells in both the apical and basolateral compartments with USUV (**Fig. 4B-C**) and WNV (**Fig. 4D-E**) via IF staining of viral envelope and respective cell markers of each compartment. All three cell types of the *in vitro* BBB were infected at 72hpi in both virus conditions. This infection did not result in significant changes in TEER compared to mock at any time-point for either virus (**Fig. 4F**). These data show that USUV invaded across the *in vitro* BBB within the same time-scale as WNV, and that neither virus induced disruption of the barrier following this initial invasion.

**Figure 4.**
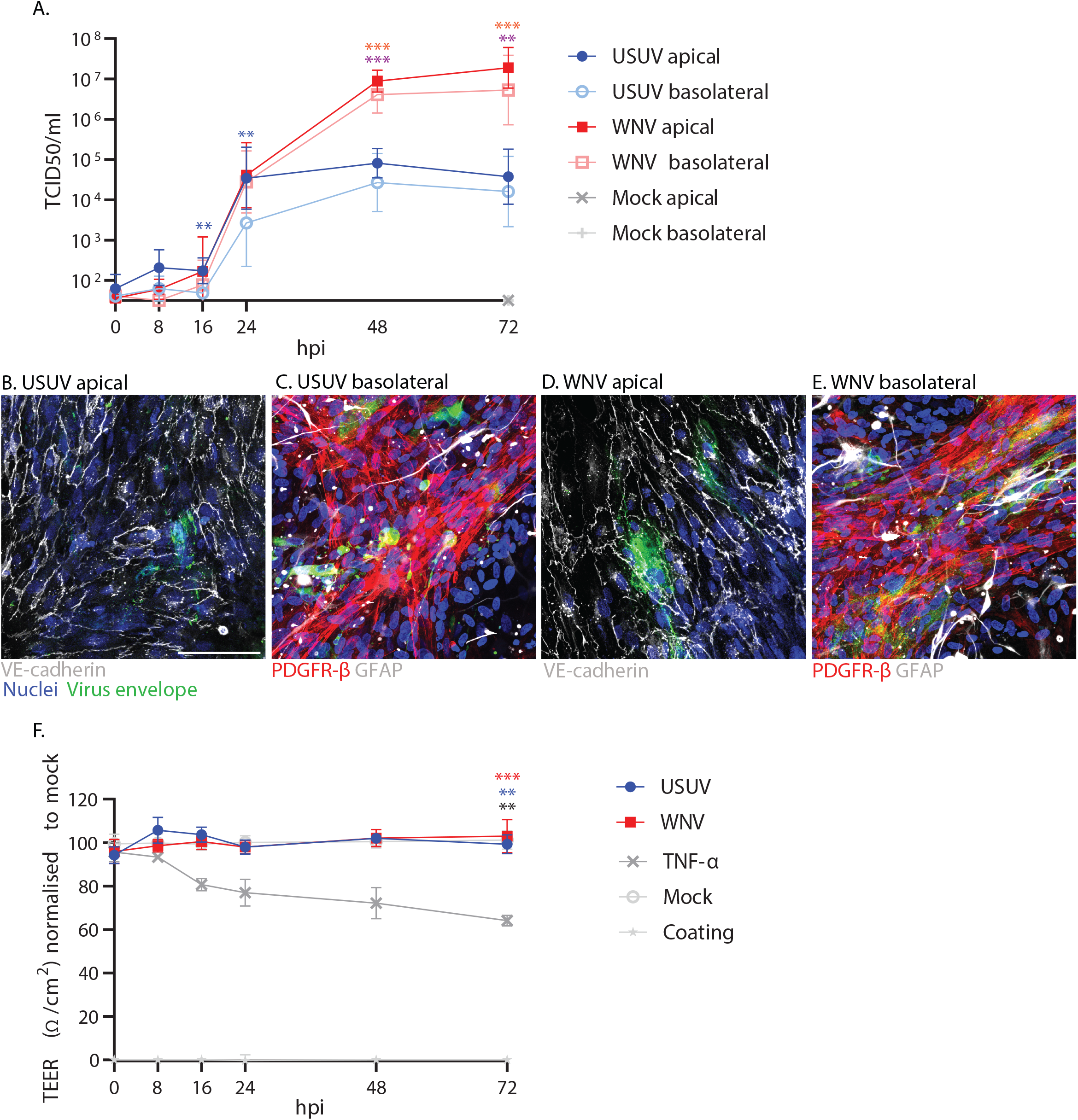
Usutu virus and West Nile virus infect and invade across the *in vitro* blood-brain barrier within the same time-frame and do not cause barrier disruption. **A.** Growth kinetics in the apical and basolateral compartments of Usutu virus (USUV) and West Nile virus (WNV) infected *in vitro* blood-brain barriers (BBB), infected at a multiplicity of infection of 1. n=2. 3 replicates per condition, per experiment. Blue stars indicate USUV apical vs USUV basolateral. 16hpi ** p=0.0089. 24hpi ** p=0.003. Purple stars indicate USUV apical vs WNV apical. *** p=0.0005. Orange stars indicate USUV basolateral vs WNV basolateral. ** p=0.0029. *** p=0.0008. Tukey’s multiple comparison. **B.** Immunofluorescent (IF) staining of USUV infected brain microvascular endothelial cells (BMECs) in apical compartment (VE-cadherin shown in white) and **C.** astrocytes and pericytes in the basolateral compartment of the *in vitro* BBB at 72hpi. Nuclei shown in blue. USUV envelope protein shown in green. PDGFR-β shown in red. GFAP shown in white. **D.** IF staining of WNV infected BMECs in apical compartment and **E.** astrocytes and pericytes in the basolateral compartment of the *in vitro* BBB at 72hpi. Representative images from 2 independent experiments. Scale bars represent 100μm. **F.** Transendothelial electrical resistance (TEER) of USUV and WNV infected and TNF-α stimulated *in vitro* BBBs across the course of the growth-kinetics experiment. n=2. 3 replicates per condition, per _26_ experiment. Red stars indicate TNF-α vs WNV. Blue stars indicate TNF-α vs USUV. Grey stars indicate TNF-α vs mock. ** p=0.0016. *** p=0.0003. Tukey’s multiple comparison.

### WNV and USUV infection of the *in vitro* BBB induces changes in key factors associated with barrier function, anti-viral immunity and immune cell transmigration

To characterise the host response mounted upon USUV and WNV infection of the *in vitro* BBB, we identified the concentration of a range of cytokines and chemokines associated with barrier disruption and immune response to viral infection in the supernatants of infected and un-infected *in vitro* BBBs (**Fig. 5A-D. Fig. S3A-L**).

**Figure 5.**
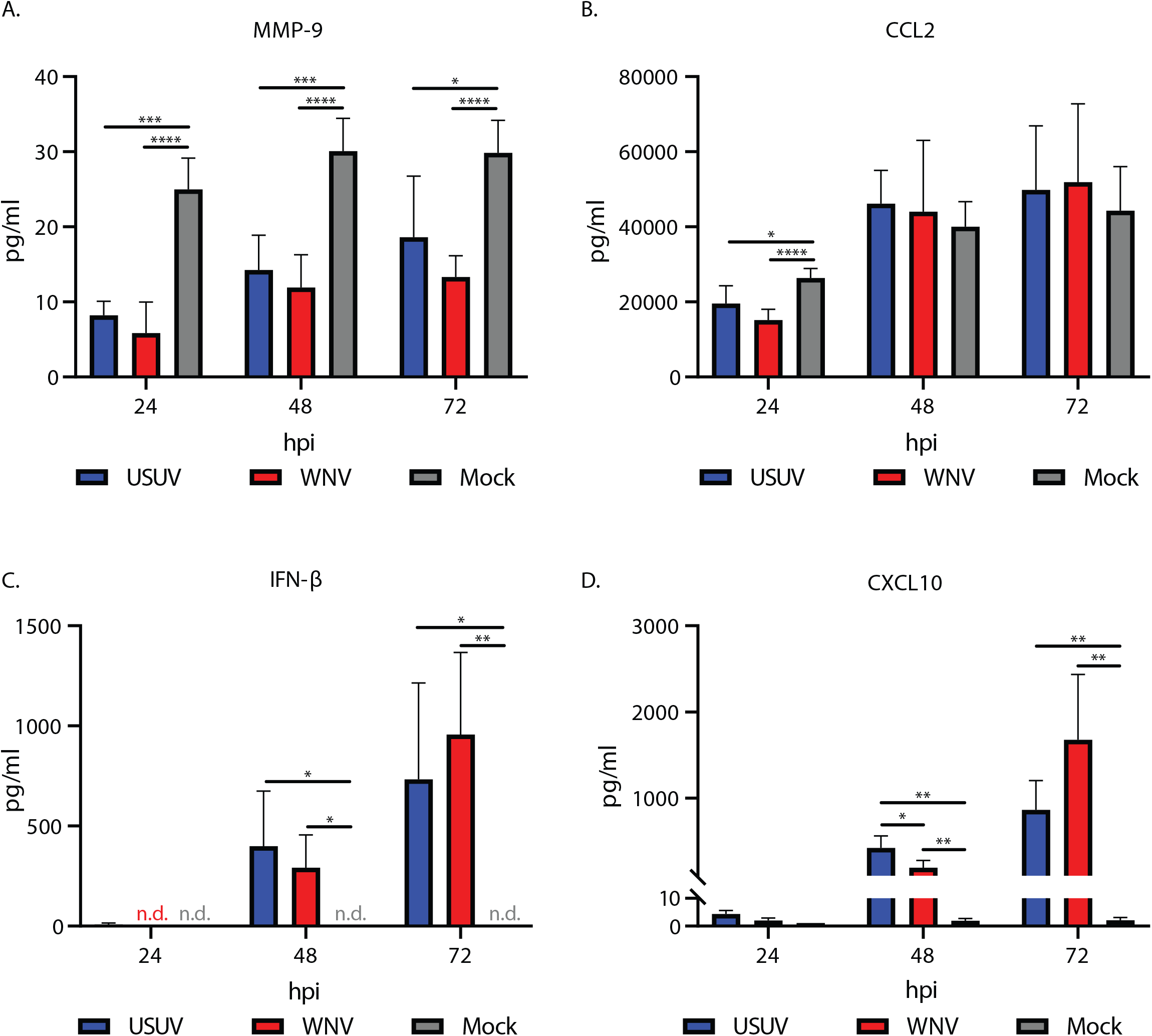
Usutu virus and West Nile virus infection of the *in vitro* blood-brain barrier induces changes in key factors associated with barrier function, anti-viral immunity and immune cell transmigration. Concentrations of **A.** MMP-9 (* p=0.0435. **** p<0.0001. 24hpi *** p=0.0001. 48hpi *** p=0.0003) **B.** CCL2 (* p=0.0368. **** p<0.0001) **C.** IFN-β (48hpi * Usutu virus (USUV) vs mock p=0.0365, West Nile virus (WNV) vs mock p=0.0162. 72hpi * p=0.0306, ** p=0.0053) and **D.** CXCL10 (* p=0.0225. 48hpi ** USUV vs mock p=0.0018, WNV vs mock p=0.006. 72hpi ** USUV vs mock p=0.0037, WNV vs mock p=0.0068.) in basolateral supernatants from *in vitro* BBBs infected at MOI 1 with WNV or USUV. Tukey’s multiple comparison. n=2. 3 replicates per condition, per experiment.

As we observed no disruption of barrier function in the infected conditions, we hypothesised that there would be little difference between infected and non-infected conditions in the expression of factors known to directly increase permeability of the BBB. USUV and WNV infected conditions had a significantly reduced concentration of MMP-9 at all time-points compared with mock (**Fig. 5A**). MMP-3 was not down-regulated in infected conditions, with both USUV and WNV having comparable concentrations to the mock-infected conditions at 24hpi and 48hpi, and increased concentrations at 72hpi (**Fig. S3J**). Additionally, we found that CCL2 was highly expressed in the *in vitro* BBB, but infection with USUV and WNV resulted in a significantly lower concentration at 24hpi compared to mock (**Fig. 5B**).

Whilst concentrations of MMP-9 and CCL2 were reduced upon infection, expression of IFN-β (**Fig. 5C**) and CXCL10 (**Fig. 5D**) was induced as the infection-course progressed, with significant increases over mock after 48hpi and 72hpi for both the USUV and WNV infected conditions. At 48hpi, USUV induced a significantly increased expression of CXCL10 compared with WNV, but this difference between viruses was no longer present at 72hpi. Overall, these data show reduction of MMP-9 and CCL2 concentrations, and an induction of IFN-β and CXCL10 across time, as a result of infection.

## DISCUSSION

WNV is thought to employ both transcellular and paracellular modes of invasion across the physical barriers that separate the blood from the CNS, such as the BBB. The BBB acts as a key entry-point for neuroinvasion, however the ability of USUV to invade the CNS across this barrier is not well understood^4^. Here we show that all three cell types of the human BBB are susceptible to infection with USUV and WNV. Previous work has shown susceptibility of human pericytes and astrocytes to infection with USUV after four days of infection^6^, but the kinetics of USUV replication in these cells, combined with a direct comparison with WNV, has not previously been carried out. We found USUV and WNV demonstrated similar replication kinetics, with comparable titres observed at 72hpi between USUV and WNV in all three cell types, despite WNV peaking higher and later than USUV in BMECs and astrocytes. This contrasts with the data obtained from our *in vitro* BBB, in which WNV grew to significantly higher titres than USUV after 24hpi. Prior to 24hpi, the kinetics of invasion were comparable between USUV and WNV, and the absence of barrier disruption at any time-point indicates a transcellular mode of invasion. A previously published *in vitro* human umbilical vein endothelial cell model, co-cultured with bovine pericytes, showed similar results for USUV^7^, however WNV was found to induce severe endothelial barrier impairment and replicated less efficiently than USUV ^8^, thereby contrasting with our data. One important difference between these previous data and our model is our use of primary human BMECs, astrocytes and pericytes in a physically proximate triple-coculture. Additionally, we primarily focused on the early-phase of invasion across the BBB, between 0 and 72hpi, not later stage effects of viral replication, up to 10dpi as previously studied^8^. Our *in vitro* data is in line with previously published *in vivo* data, describing an initial entry of WNV into the brain in absence of BBB disruption^9^ followed by a later disruption due to infection of neural cells, leading to infiltration of immune cells that contribute to barrier disruption via release of chemotactic and inflammatory factors^4,10,11^.

Previously, elevated levels of MMP-9 has been implicated in contributing to BBB disruption during WNV infection^12,13^, but in our *in vitro* BBB we observed a significant decrease in MMP-9 concentration in both USUV and WNV infected conditions. CCL2 has also been found to increase endothelial permeability^14,15^, however we observed a reduction in the concentration of CCL2 in the USUV and WNV infected conditions compared to mock at 24hpi. As both WNV and USUV infected *in vitro* BBBs showed comparable measures of barrier function to mock, the reduction in MMP-9 and CCL2 concentrations was therefore not sufficient to induce a direct, detectable decrease in permeability of the barrier as a response to infection. Increased levels of CCL2 and MMP-9 have also been associated with aiding in influx of host immune cells^16,17^. The implications and underlying mechanisms of our observed reduction of MMP-9 and CCL2 upon USUV and WNV infection require further elucidation.

Future work with the *in vitro* BBB model employed in this study should include addition of *in vitro* or *ex vivo* brain cultures in the basolateral compartment, thereby modelling the entire process of neuroinvasion and subsequent infection of the brain, and increasing the relevance of studying secondary neuroinvasion and barrier disruption of the BBB at late timepoints.

Reproducible experimental USUV infection of the CNS can only be achieved *in vivo* using neonatal mouse models, in which the BBB is not yet fully formed^18^, or in severely immune-compromised mouse models^19^. Contrastingly, WNV can induce neurological disease in immunocompetent mice^4,20^, indicating that physical or immune barriers prevent neuroinvasion of USUV. In the *in vitro* BBB, WNV reached significantly higher titres in both compartments compared to USUV. These data suggest a more effective control of USUV replication at early timepoints by the anti-viral response. USUV has previously been found to have an increased sensitivity to type I and type III interferon (IFN) responses, compared with WNV^21^, and the resistance of WNV to type I IFN signalling was found to be a key determinant in its replication fitness and virulence^22^. We saw few significant differences between USUV and WNV in the cytokine profiles of their respective basolateral supernatants, despite WNV growing to significantly higher titres, suggesting a dampened response to infection with WNV. As the BMECs in the apical compartment did not release virus into the basolateral compartment until around 16hpi, the exposure of the astrocytes and pericytes in the basolateral compartment to virus in the supernatant was delayed until this point. This may explain the relatively low presence of some of the cytokines we investigated at 24hpi, including IFN-β and CXCL10, that may be dependent on infection or activation of astrocytes or pericytes^4,23,24^.

Ultimately, the anti-viral response was activated in the infected conditions, with increasing IFN-β concentrations following the 24hr time-point. CXCL10 is an IFN-induced chemokine^25^ that promotes extravasation and activation of CXCR3 positive immune cells^26^, and has been shown to play an important role in protection of the CNS during WNV infection^27,28^. In our USUV and WNV infected *in vitro* BBBs, we saw significant induction of CXCL10 in line with increasing IFN-β concentrations. Homing of immune cells towards the site of an infection plays a paradoxical role in neuroinvasive disease, on one hand, aiding in immune-control and clearance of the virus, on the other, contributing to immunopathology and potentially exacerbating viral invasion via the Trojan horse mechanism. Invasion via Trojan horse has already been suggested for WNV^4^, however this has not been well studied *in vitro* and the potential for USUV to employ this route is unknown. Monocytes and dendritic cells have been shown to be permissive to USUV infection, and showed increased binding to an uninfected *in vitro* ‘brain-like’ endothelial barrier model, whereas non-infected immune cells showed increased binding to an infected barrier at 7dpi^8^. As we have identified that infection with USUV and WNV leads to early modulation of factors associated with attraction toward and migration across the BBB by host-immune cells, follow-up studies will focus on combination of our *in vitro* BBB with primary human immune cells to further investigate the mechanisms of (infected) leukocyte extravasation across an (infected) human BBB.

## CONCLUSIONS

In this study we aimed to identify the capacity of USUV to employ haematogenous routes of neuroinvasion and compare with WNV to gain an insight into potential mechanisms underlying the differing clinical presentation and severity resulting from infection with these viruses. We have shown that both USUV and WNV can use a transcellular mode of invasion in absence of barrier disruption. The attenuation in replicative capacity of USUV within the BBB may contribute to the observed reduction in severity and frequency of human disease, compared with WNV. However, we have shown that, when given access to the BBB, USUV is able to invade. Further work must be done to investigate the capacity of USUV to employ alternate routes of neuroinvasion and identify key factors underlying the disparate disease severity resulting from infection with USUV versus WNV.

## MATERIALS AND METHODS

### Virus strains and culturing

All viruses were grown and passaged on Vero cells (African green monkey kidney epithelial cells, ATCC CCL-81). Cells were infected at a multiplicity of infection (MOI) of 0.01 and incubated at 37 °C for 5-6 days in Dulbecco’s modified Eagle’s medium (DMEM; Lonza) with 2% FBS (Sigma-Aldrich), 100U/ml penicillin, 100μg/ml streptomycin (Lonza), 1% sodium bicarbonate (Lonza) and 2mM L-glutamine (Lonza). Supernatant was harvested, spun down at 4,000 g for 10 minutes (mins) and aliquoted then frozen at −80°C. The virus strains used in this study were selected based upon their circulation in the Netherlands and included: USUV (lineage Africa 3, GenBank accession MH891847.1, EVAg 011V-02153, isolated in 2016 from *Turdus merula*) and WNV (lineage 2, GenBank accession OP762595.1, EVAg 010V-04311, isolated in 2020 from *Phylloscopus collybita*).

### Cell culture

For human astrocytes (HA, Sciencell) and human pericytes (HP, Sciencell), flasks were coated with 2μg/cm^2^ poly-L-Lysine (Sciencell) diluted in de-ionised, sterile water and incubated at 37°C, 5% CO_2_ for a minimum of 1 hour (hr) to a maximum of 24hrs. Flasks were washed twice with de-ionised water. For human brain microvascular endothelial cells (BMECs, Cell systems) flasks were coated with 2-5ml of pre-warmed 1% gelatin (Sigma Aldrich) dissolved in PBS and incubated at 37°C, 5% CO_2_ for a minimum of 15mins to a maximum of 24hrs.

Sub-confluent cell flasks were washed with PBS before addition of 0.25% Trypsin-EDTA (Gibco) and incubation at 37°C, 5% CO_2_ for 2-5mins. Trypsin was inactivated with 5ml of FBS and 5ml of cell-type specific culture medium (Astrocyte medium; AM, Sciencell. Pericyte medium; PM, Sciencell. BMEC medium; MV2, Promocell). Cell suspensions were spun at 120g for 5mins and resuspended in their respective medium prior to passage into a pre-coated flask. HP were used between passages 3 and 10. HA were used between passages 4 and 10. BMECs were used between passages 7 and 12.

### Transwell seeding

ThinCert™ (Greiner) 12-well, translucent inserts with 3µm pores were used throughout the study. On day 1, the basolateral sides of the membrane inserts were coated by turning inserts upside-down so the basolateral side faced upward. 100μl of 2μg/cm^2^ poly-L-Lysine was pipetted onto the basolateral membrane and spread evenly with the side of a 200μl pipette tip, before incubating for a minimum of 1hr at 37°C, 5% CO_2_. Coating solution was then removed and the plate containing inserts was returned to the normal orientation. The inserts were washed twice with de-ionised water and then were inverted and left to dry in the biosafety cabinet for 1hr. 100μl of 3×10^6^/ml HA cell suspension was pipetted onto the basolateral membrane and incubated for 2-3hrs at 37°C, 5% CO_2_. Transwells were reverted and 1100μl of HA medium was added to the basolateral and 800μl was added to the apical compartment. On day 2, all medium was removed and transwells were inverted. 100μl of 6×10^5^/ml HP cell suspension was pipetted onto the basolateral membrane and incubated for 2-3hrs at 37°C, 5% CO_2_. The transwells were then reverted and 1100μl of a 1:1 mix of HA and HP medium was added to the basolateral and 800μl was added to the apical compartment. On day 4, all medium was removed from the apical compartment and replaced with 150μl of 1% gelatin solution and then incubated for 1hr at 37°C, 5% CO_2_. Gelatin solution was removed and 150μl of 7.5×10^5^/ml BMEC cell suspension was pipetted into the apical compartment and incubated for 4-6hrs at 37°C, 5% CO_2_. The basolateral medium was then replaced with 1200μl of a 1:1 mix of HA and HP medium and 800μl of MV2 medium was added to the apical compartment.

### Transendothelial electrical resistance assay

An EVOM2 trans-endothelial electrical resistance (TEER) meter with STX2 chopstick electrodes (World Precision instruments) was used to assess barrier function. TEER of a poly-L-lysine and gelatin coated transwell was used as a coating only control. Electrodes were first sterilised in 70% ethanol for 5mins, then moved to deionised water, PBS and finally a 1:1 mix of AM/PM medium. Electrodes were then placed into the apical and basolateral compartments of the transwell inserts. TEER was measured and normalised to the coating only control and surface area of the transwell to obtain normalised Ω/cm^2^.

### Infection of *in vitro* blood-brain barrier cells

For infection of the monocultured cell types, 48-well plates were coated for 1hr with 2μg/cm^2^ poly-L-Lysine, or for 15mins with 1% gelatin. HA (2.5×10^5^ cells/well), HP (2×10^5^ cells/well) and BMEC (1×10^5^ cells/well) were seeded in a 48-well plate in their respective medium and used for infection experiments the following day. *In vitro* BBB cultures were used for infection experiments after 4 days post-BMEC seeding. Barrier function of the BBB cultures was confirmed via TEER prior to infection. For infection of BBB cultures, all medium in the apical compartment was removed before addition of virus inoculum diluted in MV2 medium to an MOI of 1, calculated based on the number of BMECs. MV2 medium alone was used for the mock infected condition. For infection of monocultures, virus inoculum diluted to an MOI of 1 in the cell-type specific medium was used for infection. Plates were returned to the incubator at 37°C for 1hr then virus inoculum was removed and the well, in case of monoculture infection, or apical compartment, in case of BBB culture infection, was washed three times with PBS before addition of cell-type specific medium. For the BBB cultures, sample was taken from both the apical and basolateral compartments for time-point 0hpi, and transwells were then transferred to a clean plate with fresh AM/PM basolateral medium. Plates were returned to the incubator at 37°C. When stated, as a positive control for barrier disruption, uninfected cultures were treated with 100ng/ml of TNF-α in the apical compartment, which was maintained throughout the experimental course. Supernatant was removed and refreshed at the specified timepoints. The harvested supernatants were stored at −80°C for titration or RNA isolation.

### Virus titration

Tenfold serial dilutions of culture supernatants were inoculated onto a semiconfluent monolayer of Vero cells in a 96-well plate (2.3 × 10^4^ cells/well). Cytopathic effect (CPE) was used as titre read out and determined at 6 days post-infection (dpi). Virus titres were calculated as the 50% tissue culture infective dose (TCID50) using the Spearman-Kärber method^29^. An initial 1:10 dilution of supernatant resulted in a detection limit of 31.6 TCID50/ml.

### RNA isolation and real-time quantitative PCR for quantification of virus

Sample supernatants in MagnaPure lysis buffer were incubated with Agencourt AMPure-XP (Beckman Coulter) magnetic beads in a 96-well plate. The plate was placed on a DynaMagTM-96 magnetic block (Invitrogen) and supernatant was removed. The beads were washed three times with 70% ethanol whilst on the magnetic block and then left to air dry. The plate was removed from the block and beads were resuspended in de-ionised water to elute the isolated RNA.

A real-time TaqManTM assay was performed using the Applied-Biosystems 7500 real-time PCR system (ThermoFisher Scientific). WNV primer/probe mix (Forward-primer sequence: CCACCGGAAGTTGAGTAGACG, Reverse-primer sequence: TTTGGTCACCCAGTCCTCCT, Probe sequence: TGCTGCTGCCTGCGGCTCAACCC) was diluted in TaqManTM fast virus 1-Step Master Mix and de-ionised water to a final volume of 12µl before addition of 8µl sample RNA. The following program was used: 5 min 50 °C, 20 sec 95 °C and 45 cycles of 3 sec 95 °C and 30 sec 60 °C. Samples were compared to a standard-curve of virus stock dilutions to acquire a TCID50 equivalent/ml.

### Dextran permeability assay

20kDa fluorescent TRITC dextran (Sigma Aldrich) was added to the apical compartment of the *in vitro* BBB at 12hpi to a final concentration of 100µg/ml, and incubated until 24hpi. Fluorescence of the basolateral supernatant was assessed using a Tecan Infinite F200 Pro. Data was normalised to fluorescence in the mock infected condition. BBB cultures treated with 100ng/ml of TNF-α were used as a positive control for barrier disruption.

### Multiplex Immunoassay

To determine concentrations of a panel of human cytokines and chemokines in cell culture supernatants, a custom human magnetic Luminex screening assay, 16-plex kit (R&D systems) was used according to the manufacturer’s instructions. Measured proteins were: CCL2, CXCL10, G-CSF, GM-CSF, ICAM-1, IFN-α, IFN-β, IL-1β, IL-6, IL-8, IFN-lambda 2, IFN-lambda 3, MMP-3, MMP-9, TNF-α and VCAM-1. Samples were read using a Bio-Plex suspension array system with automated concentration calculation based upon relation to standard curves.

### Immunofluorescent staining

Cells were fixed for 30mins in 10% formalin. Fixed cells were permeabilised with 0.5% triton (Sigma) diluted in PBS for 15mins, then blocked with 5% bovine serum albumin (BSA; Aurion) for 1hr before incubation with primary-antibodies diluted in PBS with 2% BSA overnight at 4°C. Cells were washed three times in PBS then incubated with secondary-antibodies for 1hr at room temperature (RT) in the dark. Cells were washed then incubated with Hoechst (1:1000, Invitrogen) for 20mins at RT in the dark. Images were obtained using a Zeiss LSM 700 laser scanning microscope. Primary-antibodies used in this study were: mouse anti-flavivirus envelope protein (1:250, D1-4G2-4-15 hybridoma; ATCC, USA), goat anti-VE-Cadherin (1:100, R&D systems), rabbit anti-PDGFRβ (1:200, clone Y92, Abcam), rabbit anti-GFAP (1:200, Millipore), chicken anti-GFAP (1:200, Abcam). An AF555 conjugated mouse antibody was used to stain ZO-1 (1:100, Thermofisher Scientific) followed by incubation with a secondary-antibody to amplify the fluorescent signal. The secondary-antibodies (Invitrogen) used in this study were: anti-mouse AF488 (1:250), anti-rabbit AF488 (1:200), anti-mouse AF555 (1:200), anti-rabbit AF555 (1:200), anti-chicken AF555 (1:200), anti-rabbit AF647 (1:200), anti-chicken AF647 (1:200) and anti-goat AF647 (1:200), hosted in donkey.

### Image processing

Images underwent processing and file-type conversion using ImageJ software (version 1.53t, National Institutes of Health, Bethesda, MD). Processed images were rendered in 3D using Dragonfly software (Version 2021.1 for [Windows]; Comet Technologies Canada Inc., Montreal, Canada). This software is available at https://www.theobjects.com/dragonfly.

### Statistical analysis

Quantitative data were analysed and statistics carried out using Prism 8.0.2 (GraphPad).

## DATA AVAILABILITY

The datasets used and/or analysed during the current study available from the corresponding author on reasonable request.

## ACKNOWLEDGEMENTS

We thank Danny Noack (Erasmus MC) for his useful discussions and technical help running the multiplex immunoassay.

---

**Supplementary figure 1.**
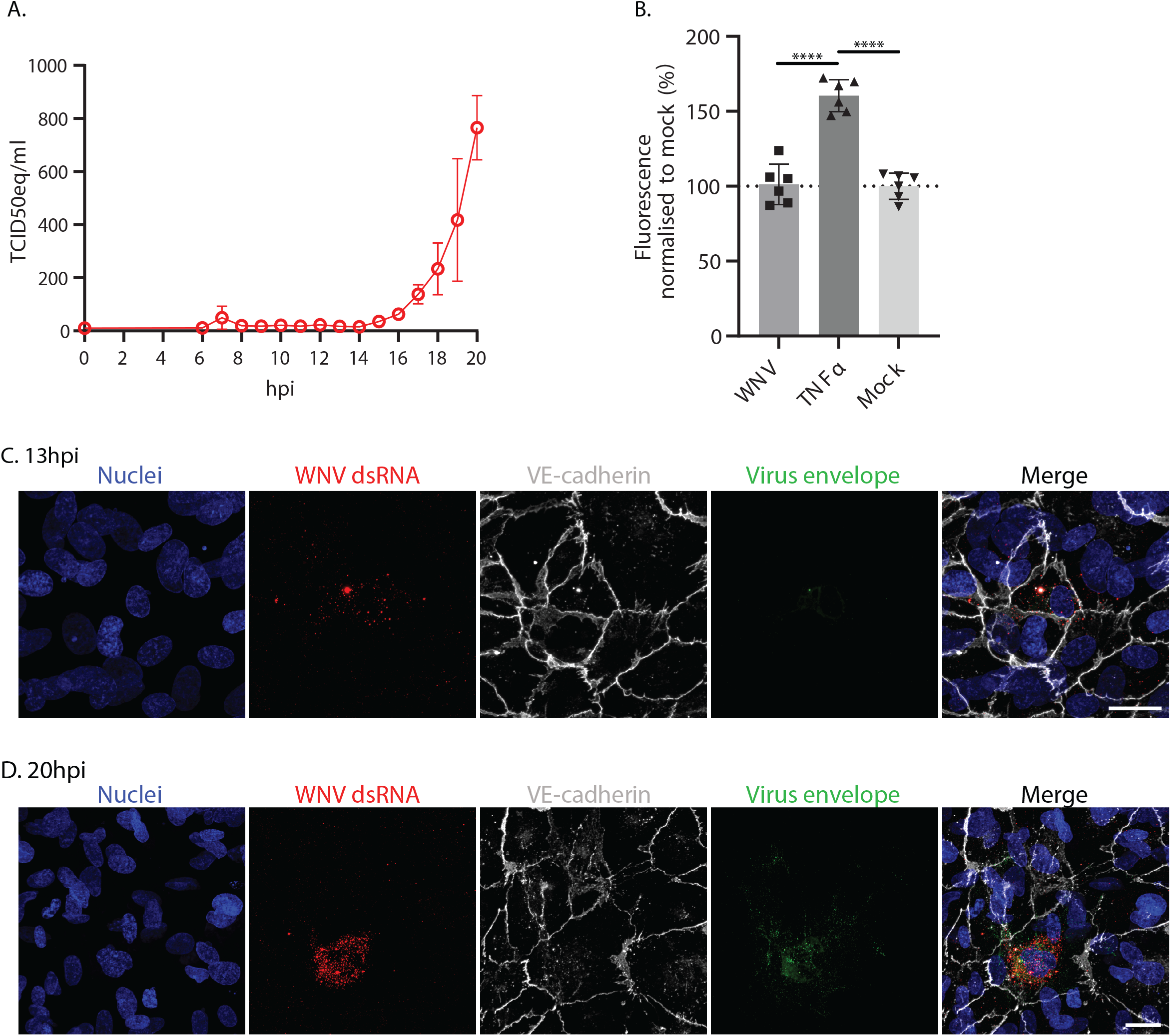
West Nile virus invades across the *in vitro* blood-brain barrier in absence of barrier disruption, after an initial round of replication in brain microvascular endothelial cells. **A.** TCID50 equivalents of West Nile virus (WNV) in the basolateral compartment of the *in vitro* blood-brain barrier (BBB) infected at a multiplicity of infection of 1. Two replicates were used per time-point, for a total of 32 *in vitro* BBBs across the course of the experiment. Representative data from 2 independent experiments. qPCR was used to quantify presence of viral genome in cell lysates which was compared against a standard curve of diluted virus stock to obtain TCID50eq values. B. Leakage of a 20kDa fluorescent dextran through the *in vitro* BBB model at 24hpi with WNV at an MOI of 1 or stimulation with TNF-α, relative to mock infection. **** p<0.0001. One-way ANOVA. C. IF staining of WNV infected BMECs in apical compartment of *in vitro* BBB at 13hpi and D. 20hpi. Scale bars represent 50μm.

**Supplementary figure 2.**
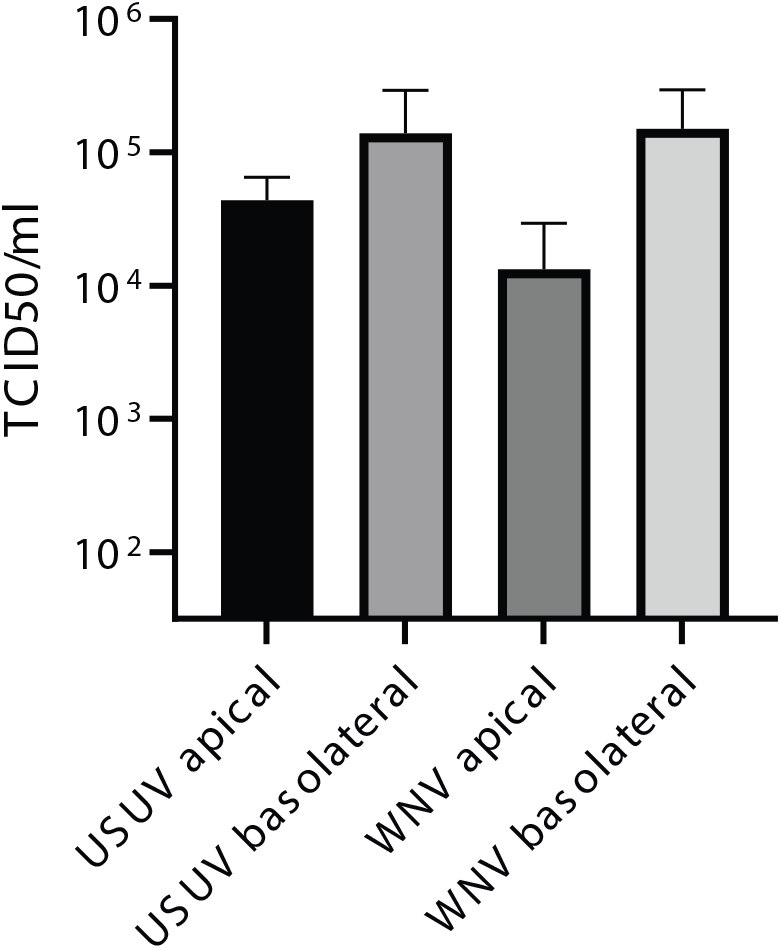
The transwell membrane does not prevent diffusion of virus into the basolateral compartment. Titres of Usutu virus and West Nile virus in apical and basolateral compartments of empty, coated transwells immediately following the inoculation incubation. n=1. 3 replicates.

**Supplementary figure 3.**
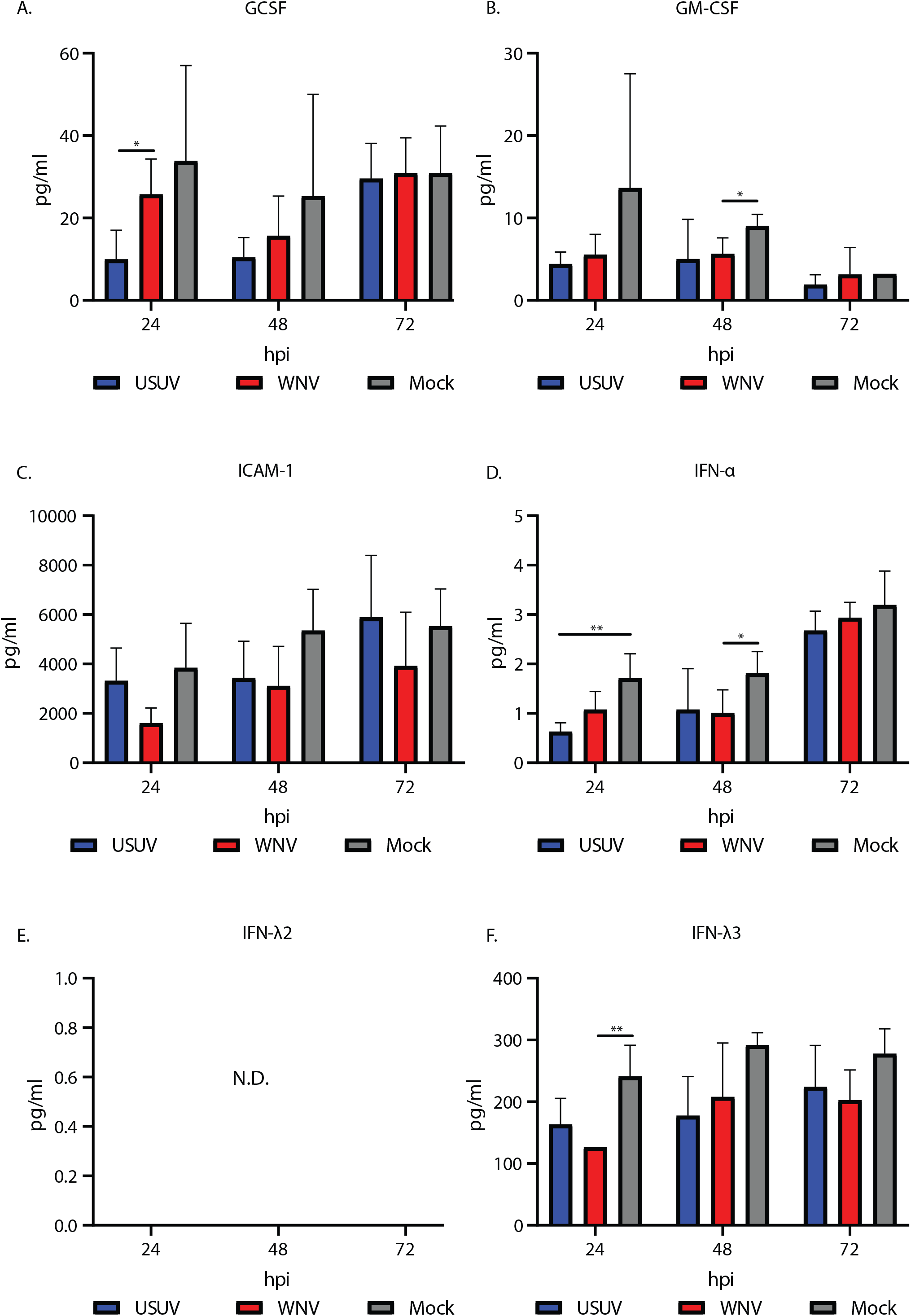
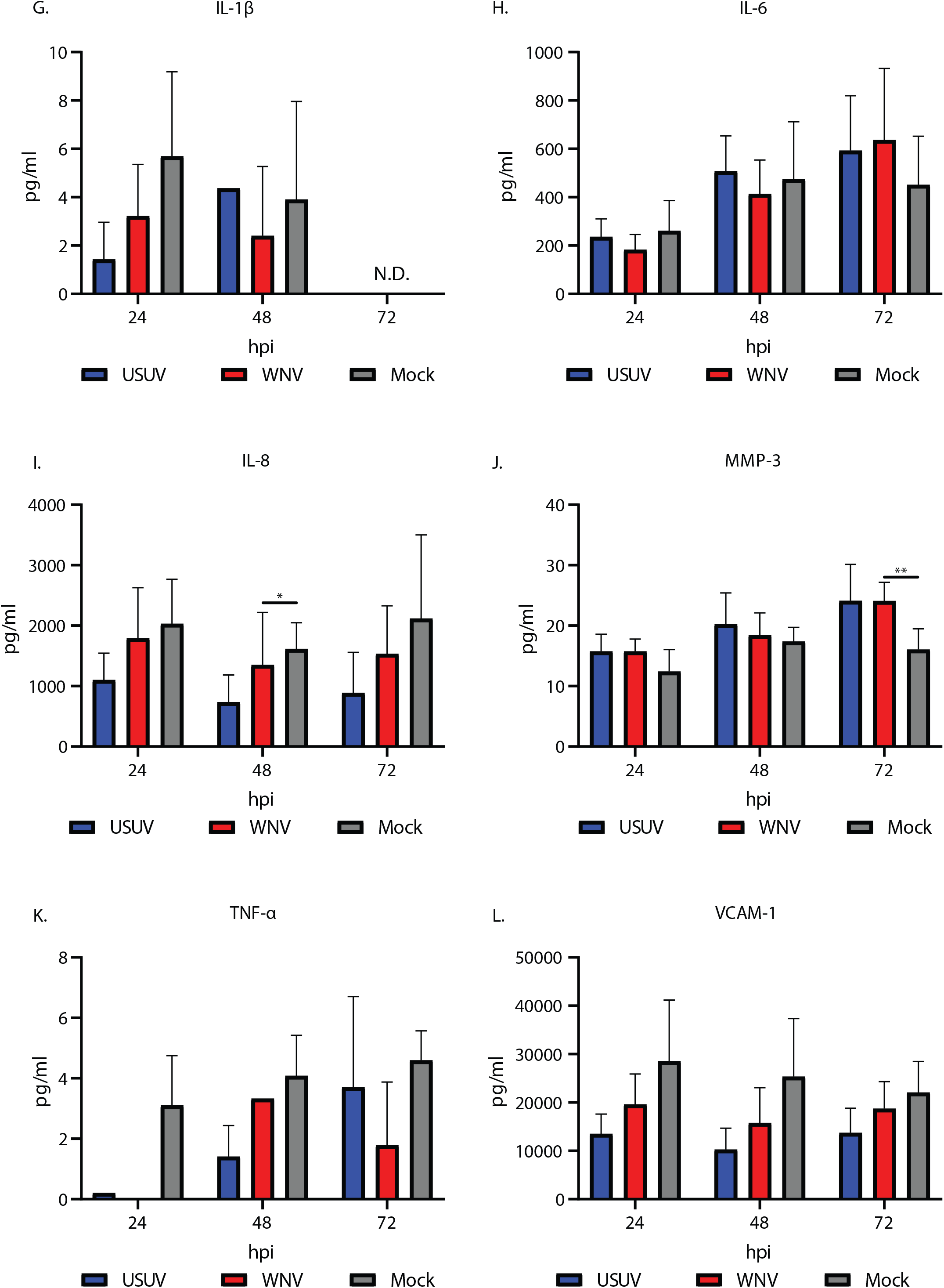
Concentration of **A.** G-CSF (* p=0.0214), **B.** GM-CSF (*p=0.0176), **C.** ICAM-1, **D.** IFN-α (* p=0.0295. ** p=0.0048), **E.** IFN-λ2, **F.** IFN-λ3 (** p=0.0057), **G.** IL-1β, **H.** IL-6, **I.** IL-8 (* p=0.0238), **J.** MMP-3 (** p=0.0043), **K.** TNF-α, **L.** VCAM-1 in basolateral supernatants from *in vitro* blood-brain barriers infected at a multiplicity of infection of 1 with West Nile virus (WNV) or Usutu virus (USUV). Tukey’s multiple comparison. n=2. 3 replicates per condition, per experiment.

**Supplementary video 1.** 3D rendering of the triple-cocultured *in vitro* blood-brain barrier (BBB) model layout with brain microvascular endothelial cells (BMECs) in apical compartment, astrocytes and pericytes in basolateral compartment and the transwell membrane separating the two. VE-cadherin shown in white. GFAP shown in red. PDGFR-β shown in green. Nuclei shown in blue. Dragonfly ORS used for rendering of confocal images.

## REFERENCES

1. Historic Data (1999-2022) | West Nile Virus | CDC. https://www.cdc.gov/westnile/statsmaps/historic-data.html.

2. Historical data by year - West Nile virus seasonal surveillance. https://www.ecdc.europa.eu/en/west-nile-fever/surveillance-and-disease-data/historical.

3. Angeloni, G. et al. Epidemiology, surveillance and diagnosis of Usutu virus infection in the EU/EEA, 2012 to 2021. Euro Surveill 28, 2200929 (2023).

4. Marshall, E. M., Koopmans, M. P. G. & Rockx, B. A Journey to the Central Nervous System: Routes of Flaviviral Neuroinvasion in Human Disease. Viruses 14, 2096 (2022).

5. Stone, N. L., England, T. J. & O’Sullivan, S. E. A novel transwell blood brain barrier model using primary human cells. Front Cell Neurosci 13, 455689 (2019).

6. Clé, M. et al. Differential neurovirulence of Usutu virus lineages in mice and neuronal cells. J Neuroinflammation 18, 1–22 (2021).

7. Clé, M. et al. Study of usutu virus neuropathogenicity in mice and human cellular models. PLoS Negl Trop Dis 14, 1–27 (2020).

8. Constant, O. et al. Differential effects of Usutu and West Nile viruses on neuroinflammation, immune cell recruitment and blood–brain barrier integrity. Emerg Microbes Infect 12, (2023).

9. Roe, K. et al. West nile virus-induced disruption of the blood-brain barrier in mice is characterized by the degradation of the junctional complex proteins and increase in multiple matrix metalloproteinases. Journal of General Virology 93, 1193–1203 (2012).

10. Salinas, S. et al. Deleterious effect of Usutu virus on human neural cells. PLoS Negl Trop Dis 11, e0005913 (2017).

11. Kumar, M., Verma, S. & Nerurkar, V. R. Pro-inflammatory cytokines derived from West Nile virus (WNV)-infected SK-N-SH cells mediate neuroinflammatory markers and neuronal death. J Neuroinflammation 7, 73 (2010).

12. Verma, S., Kumar, M., Gurjav, U., Lum, S. & Nerurkar, V. R. Reversal of West Nile virus-induced blood-brain barrier disruption and tight junction proteins degradation by matrix metalloproteinases inhibitor. Virology 397, 130–138 (2010).

13. Wang, P. et al. Matrix Metalloproteinase 9 Facilitates West Nile Virus Entry into the Brain. J Virol 82, 8978–8985 (2008).

14. Bose, S. & Cho, J. Role of chemokine CCL2 and its receptor CCR2 in neurodegenerative diseases. Arch Pharm Res 36, 1039–1050 (2013).

15. Roblek, M. et al. CCL2 is a vascular permeability factor inducing CCR2-dependent endothelial retraction during lung metastasis. Mol Cancer Res 17, 783 (2019).

16. Elkington, P. T. G., O’kane, C. M. & Friedland, J. S. The paradox of matrix metalloproteinases in infectious disease. Clin Exp Immunol 142, 12–20 (2005).

17. Cabral-Pacheco, G. A. et al. The Roles of Matrix Metalloproteinases and Their Inhibitors in Human Diseases. Int J Mol Sci 21, 9739 (2020).

18. Weissenböck, H., Bakonyi, T., Chvala, S. & Nowotny, N. Experimental Usutu virus infection of suckling mice causes neuronal and glial cell apoptosis and demyelination. Acta Neuropathol 108, 453–460 (2004).

19. Benzarti, E. et al. New insights into the susceptibility of immunocompetent mice to Usutu virus. Viruses 12, (2020).

20. Graham, J. B., Swarts, J. L. & Lund, J. M. A Mouse Model of West Nile Virus Infection. Curr Protoc Mouse Biol 7, 221 (2017).

21. Cacciotti, G. et al. Variation in interferon sensitivity and induction between Usutu and West Nile (lineages 1 and 2) viruses. Virology 485, 189–198 (2015).

22. Keller, B. C. et al. Resistance to Alpha/Beta Interferon Is a Determinant of West Nile Virus Replication Fitness and Virulence. J Virol 80, 9424–9434 (2006).

23. Jansson, D. et al. A role for human brain pericytes in neuroinflammation. J Neuroinflammation 11, 1–20 (2014).

24. Abbott, N. J., Rönnbäck, L. & Hansson, E. Astrocyte-endothelial interactions at the blood-brain barrier. Nat Rev Neurosci 7, 41–53 (2006).

25. Buttmann, M., Berberich-Siebelt, F., Serfling, E. & Rieckmann, P. Interferon-β Is a Potent Inducer of Interferon Regulatory Factor-1/2-Dependent IP-10/CXCL10 Expression in Primary Human Endothelial Cells. J Vasc Res 44, 51–60 (2007).

26. Prizant, H. et al. CXCL10 + peripheral activation niches couple preferred sites of Th1 entry with optimal APC encounter. Cell Rep 36, (2021).

27. Klein, R. S. et al. Neuronal CXCL10 Directs CD8+ T-Cell Recruitment and Control of West Nile Virus Encephalitis. Journal of virology 79, 11457–11466 (2005).

28. Zhang, B., Chan, Y. K., Lu, B., Diamond, M. S. & Klein, R. S. CXCR3 Mediates Region-Specific Antiviral T Cell Trafficking within the Central Nervous System during West Nile Virus Encephalitis. The Journal of Immunology 180, 2641–2649 (2008).

29. Kärber, G. Beitrag zur kollektiven Behandlung pharmakologischer Reihenversuche. Naunyn Schmiedebergs Arch Exp Pathol Pharmakol 162, 480–483 (1931).

